# Circulating immune cell populations at rest and in response to acute endurance exercise in young adults with cerebral palsy

**DOI:** 10.1101/2023.03.22.532927

**Authors:** Annika Kruse, Ian Imery, Linnéa Corell, Emma Hjalmarsson, Rodrigo Fernandez-Gonzalo, Ferdinand von Walden, Stefan M. Reitzner

**Author notes:** these authors contributed equally (Annika Kruse, Ian Imery, Ferdinand von Walden, Stefan M. Reitzner). Corresponding author: Dr. Annika Kruse, Institute of Human Movement Science, Sport and Health, University of Graz, Mozartgasse14, 8010 Graz, Austria.

## Abstract

**Aim:** Low physical activity alters immune function and increases the risk of developing chronic inflammation. This cross-sectional study aimed at determining the immune status and function in young adults with cerebral palsy (CP) in comparison to typically developing (TD) individuals.

**Method:** Blood samples from 12 individuals with CP and 17 TD were collected before, immediately after, and one hour following 45 minutes of Frame Running or running, respectively. Independent t-tests were used to compare heart rate, level of exertion, and baseline cell proportions between groups. Mixed model ANOVA was utilized to investigate immune cell responses to exercise across groups.

**Results:** Baseline levels of TCRγδ+ T-cells were significantly higher in the individuals with CP. Several cell populations showed significant changes after exercise in both CP and TD groups. CD8+ T-cells were only significantly elevated immediately after exercise in the TD participants. Individuals with CP exhibited significantly lower heart rates, despite similar ratings of perceived exertion.

**Interpretation:** Elevated baseline TCRγδ+ T-cells may indicate low-grade inflammation in adults with CP. Although most of the cell populations showed typical responses to endurance exercise, the absence of response in CD8+ T-cells in individuals with CP may indicate the need for higher intensity during exercise.

**What this paper adds:**

- This is the first study addressing immune cells in adults with CP
- TCRγδ+ T-cell baseline levels are elevated in adults with CP
- The CD8+ T-cell response to exercise was blunted in adults with CP
- Exercise intensity is decisive for CD8+ T-cell responses in individuals with CP

## Evidence of altered immune function in individuals with CP

Cerebral palsy (CP) comprises a heterogenic group of neuropediatric disorders sharing common symptoms, mainly involving difficulties with movement and posture.^1^ CP results from a non-progressive brain lesion occurring before or during early infancy. Individuals with CP present impairments such as muscle weakness, co-contraction, spasticity, and poor selective motor control as well as balance.^1^ These features in turn are considered possible causes of various musculoskeletal pathologies that further develop into bone deformities and joint instability.^2^ Moreover, previous studies showed that muscles of individuals with spastic CP are structurally and mechanically different compared to those of typically developing (TD) individuals.^2^

To date, it is well established that individuals with CP are less physically active compared to TD individuals.^3^ Individuals with CP with low motor function are the most sedentary group and usually use assistive mobility devices, which further negatively impact their activity level, cardiorespiratory fitness, and muscle health. Combining their reduced physical activity with other factors, such as nutrient deficiencies, individuals with CP have an increased risk for developing metabolic disorders such as obesity, type 2 diabetes, arthritis, and atherosclerosis.^4,5^ All of these disorders have been associated with a chronic inflammatory status in TD individuals.^6,7^ However, little is known about the existence of chronic inflammation and/or altered immune function in adults with CP.

Although studies in individuals with CP are scarce, evidence exists indicating the occurrence of systemic low-grade inflammation and immune dysregulation in young individuals with CP.^8–13^ For instance, enhanced levels of specific cytokines (e.g., tumor necrosis factor alpha (TNF-α) and Interleukin (IL)-6)^12,13^ as well as T-cell (e.g., CD3+, CD4+, and CD8+) and B-cell subsets (CD22+) were observed in young children with CP when compared to TD children.^13^ In addition significantly increased levels of erythropoietin and a hyporesponsiveness to lipopolysaccharide for IL-1a, IL-1b, IL-2, and IL-6 levels were found in school-aged children with CP,^9^ an elevated gene expression of pro-inflammatory cytokines in skeletal muscles with fixed contractures^11^ and significantly increased levels of transforming growth factor beta-1 (TGFβ1) and C-reactive protein (CRP) were also found in children and adolescents with CP.^10^ The reported results indicate the existence of a systemic low-grade inflammation and altered immune function in children with CP. However, there is a lack of knowledge about the immune status of young adults with CP. Thus, more detailed investigations of immune cell subpopulations in young adults with CP are warranted.

Exercise influences the activation state of the immune system^14^ and also affects the inflammatory status and gene expression in its target tissues, e.g., muscle tissue.^14^ In response to physical activity, the immune cells infiltrate the tissues and mediate changes in physical performance capacity by expressing cytokines, producing antibodies, or modulating inflammation, defense, and regeneration processes.^15^ Importantly, the expression pattern of these responses and their extent depend on a number of factors such as work intensity, duration and frequency, and the amount of activated muscle mass. In healthy populations, acute bouts of endurance exercise typically lead to characteristic biphasic changes in the numbers of circulating lymphocytes (i.e., T-cells and subpopulations, B-cells):^7,16^ increase in cell numbers (i.e., lymphocytosis) during and immediately after exercise and decrease in numbers below preexercise levels during the early stages of recovery. NK-cells are rapidly mobilized into the circulation during exercise and decline within a few hours afterwards.^7^ Moreover, acute endurance exercise increased the number of circulating neutrophils.^17^ While these responses have been found in typically developed populations, there are currently no studies about the immune system response to acute exercise in young adults with CP. Information about the immune system response to exercise may uncover an altered behavior of the immune system in this group. This in turn could shift our focus on the evaluation of treatment strategies that may have the potential to better stimulate health-related immune cell responses in individuals with CP.

The main aims of this study were 1) to quantify and compare the abundance and proportions of circulating immune cell populations in young adults with CP and TD adults at rest, and 2) to examine the response of circulating immune cell populations after an acute bout of endurance exercise in individuals with CP and TD individuals. We hypothesized that the resting proportions would not differ between groups and that an endurance exercise bout would lead to a similar change in circulating immune cells in the individuals with CP and TD.

## Methods

### Study design

In this cross-sectional study, both individuals with CP and TD participants performed an acute endurance exercise bout consisting of 45 min of Frame Running or running, respectively. Blood samples were taken before, immediately after, and one hour after the 45 min endurance exercise bout. The distance covered and heart rate were measured throughout the endurance exercise, while the level of perceived exertion (Borg-RPE) was assessed directly before and every 10 min during running in both groups.

### Participants and recruitment

Altogether, 12 individuals with CP and 17 TD individuals were included in this study. The adults with CP were included based on the following criteria: diagnosis of CP, between 18-40 years of age, GMFCS level II-IV, and Frame Running experience ≥ three months. TD individuals were included if they were in the age range of 18-40 years. Exclusion criteria for both groups were soft tissue surgery during the last six months or bone surgery in the last 12 months before study participation.

All experiments carried out in this study were approved by the national ethical review board (Ethical permit 2021-05116), Sweden and written informed consent was obtained from each participant in advance.

### Anthropometries and participants’ characteristics

The participants’ anthropometrics and characteristics are displayed in Table 1. For the individuals with CP, body mass was determined by use of a wide electronic scale (Detecto 6550, Carterville, USA), which allowed weighing while sitting in a wheelchair or chair. Body height was measured with a tape measure and the participants either lying on an examination bench or while standing against a wall if possible. Body mass and height were assessed in the TD participants accordingly.

**Table 1.** Participant characteristics of the individuals with cerebral palsy (CP) and the typically developed (TD) participants. Mean (Standard deviation).

|  | **CP** | **TD** |
| --- | --- | --- |
| **Number (n)** | 12 | 17 |
| **Age (years)** | 25.1 (5.9) | 31.4 (6.2) |
| **Body mass (kg)** | 54.4 (10.6) | 72.7 (14.6) |
| **Body height (cm)** | 163.4 (11.7) | 173.0 (11.6) |
| **Body mass index (kg/m<sup>2</sup>)** | 20.3 (3.1) | 24.2 (3.3) |
| **GMFCS level (II/III/IV)** | 2/3/7 | n.a. |
| **CP type (spastic/dyskinetic/ataxic)** | 4/7/1 | n.a. |
| **FR experience (years)** | 8.5 (2.6) | n.a. |
| **Distance 6-MFRT (m)** | 886.7 (281.2) | n.a. |
| **SGPALS (1/2/3/4)** | 1/3/5/3 | 1/1/11/3 |

GMFCS, Gross Motor Function Classification System; FR, Frame Running; 6-MFRT, 6-Minute Frame Running Test; SGPALS, Saltin Grimby Physical Activity Level Scale^18^ (grading: 1: Physically inactive (I); 2: Some light physical activity (LPA); 3: Regular physical activity and training (moderate PA); 4: Regular hard physical training for competition sports (vigorous PA)); n.a., not applicable.

Moreover, all participants graded their activity level by use of the Saltin Grimby Physical Activity Level Scale.^18^ The levels are graded as follows: 1: physically inactive, 2: some light physical activity, 3: regular physical activity and training, and 4: regular hard physical training for competition sports.

### Endurance exercise bout

To assess the immunological response to acute exercise, all participants performed a 45 min run on an indoor running track (200 m), with the individuals with CP using their Frame Runners. All participants were asked to run as far as they could within the given time limit, aiming for a perceived exertion level ≥ 15 on the Borg RPE scale throughout the exercise bout. Running laps were manually counted and heart rate data was continuously collected with a heart rate monitor (Garmin Edge 25, Garmin).

To evaluate running intensity, mean heart rates were calculated over the last minute of every block of 10 minutes. Moreover, exertion level was rated before the start of the bout and every 10 minutes during running using the Borg RPE scale.^19^ For the analyses, mean values of both variables were calculated.

### Blood sampling

Peripheral venous blood was taken from the *Fossa cubitalis* using a venous catheter (PVC) directly before, immediately after, and one hour after the endurance exercise. A local anesthetic cream (EMLA, Aspen Pharma Trading Limited) was placed on the skin at least 1 hour before the first blood sample collection to reduce discomfort and pain. At all three timepoints, blood was sampled using a density gradient reagent for isolation of peripheral mononuclear blood cells (BD CPT Vacutainer #362753). Peripheral mononuclear blood cells (PMBCs) were extracted by centrifugation at 1700g for 20 minutes. After extraction from whole blood, the PMBCs were washed twice and stored in a fetal bovine serum +10% DMSO cell stock solution at −200°C.

### Immune cell analysis

Following the complete collection from all participants, frozen samples were carefully thawed for analysis. Subtypes of PBMCs (see below) were quantified using fluorescence-activated cell sorting (FACS) based on combinations of cell surface markers. For FACS analysis, about 500,000 cells were taken from each previously made cell stock to perform antibody staining in two separate panels (250,000 cells each). In the myeloid panel, PBMCs were stained for CD45 (BD #555482), CD177 (BD #564239), CD66b (BD #562254), CD19 (BD #560728), CD20 (BD #563067), CD14 (BD #555399) and CD16 (BD #562874). For the lymphoid panel, PBMCs were stained for CD8 (BD #563919), CD4 (BD #561843), CD45 (BD #555482), CD3 (BD #561810), TCRγδ (BioLegend #331221), CD16 (BD #562874) and CD56 (BD #560842). These panel markers were used in a gating strategy (Figure 1) to identify eight populations of PBMCs: B-cells, Monocytes, Macrophages, bright and dim NK-cells, CD8+ T-cells, CD4+ T-cells and TCRγδ+ T-cells.

**Figure 1.**
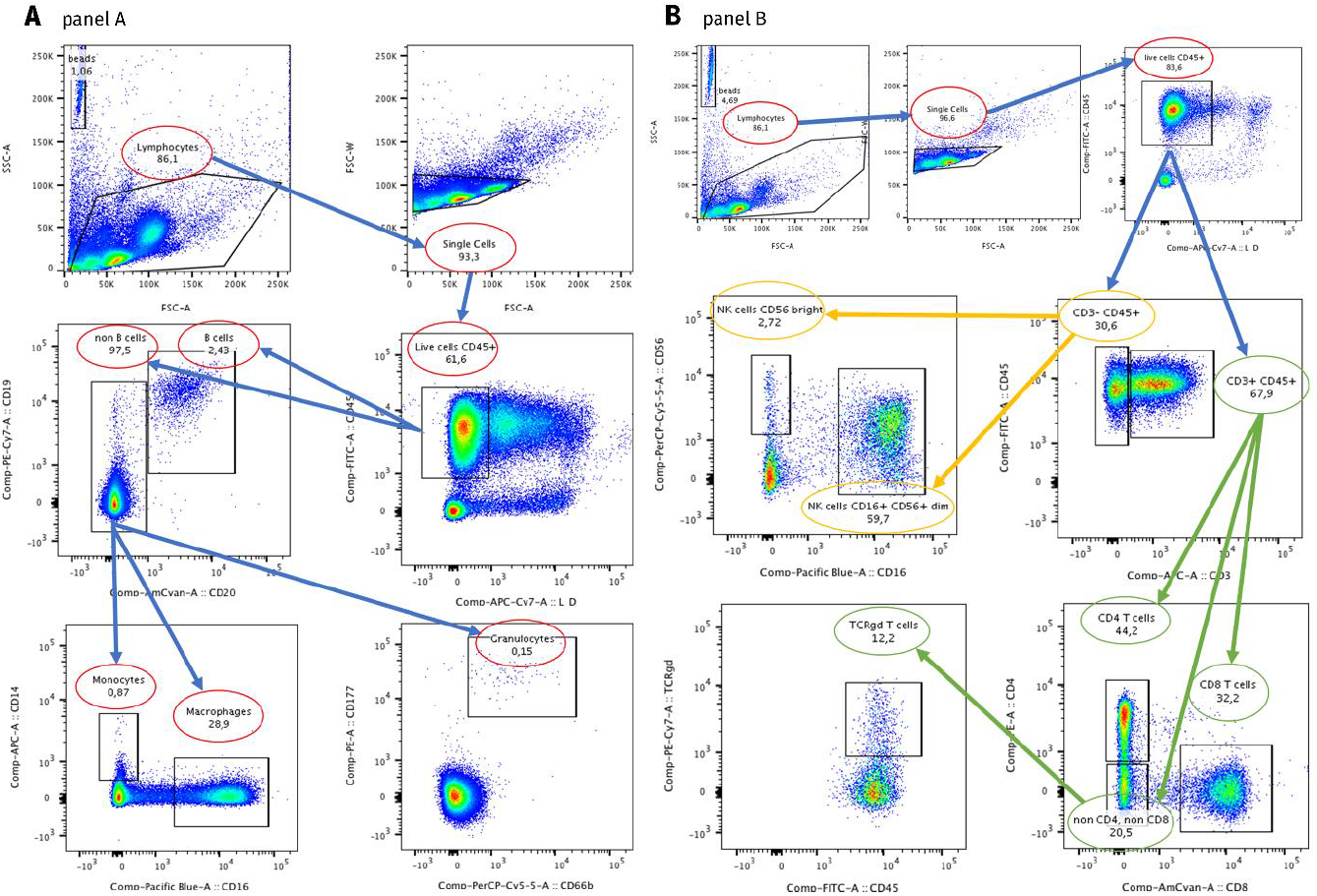
Gating strategy for the FACS analysis. Panel A (A) to identify myeloid immune cells monocyte and macrophage populations, and the lymphoid B-cells. Panel B (B) to identify lymphoid cell populations bright and dim natural killer cells, CD8+ T-cell, CD4+ T-cell and TCRγδ+ T-cell populations.

### Statistical analysis

Data was tested for normal distribution using the Shapiro-wilk test. Mean heart rates and Borg-Scale ratings were compared between the two groups (CP vs. TD) using an independent t-test (GraphPad Prism 5, GraphPad Software, San Diego, California, USA). To assess the proportions of the circulating immune cell populations (% of total cells) between groups at baseline, an independent t-test was utilized, while changes in baseline proportions were analyzed over time and between groups using a mixed model ANOVA (R, RStudio, version 3.6.0., Boston, Massachusetts, USA). Repeated testing corrections were conducted using the Tukey’s Test. Alpha was set to 0.05. Values are reported as means and standard deviations if not stated otherwise. Due to technical issues, exercise performance data of one participant had to be excluded from the data analysis.

## Results

### Exercise performance and baseline immune cell populations

Significantly lower mean heart rates (*p* < 0.05) during the 45 min running exercise were observed in the young adults with CP compared to TD adults (Figure 2A). In contrast, no difference was observed in perceived exertion (TD: 15 (1); CP: 15 (1);*p* > 0.05, Figure 2B).

**Figure 2.**
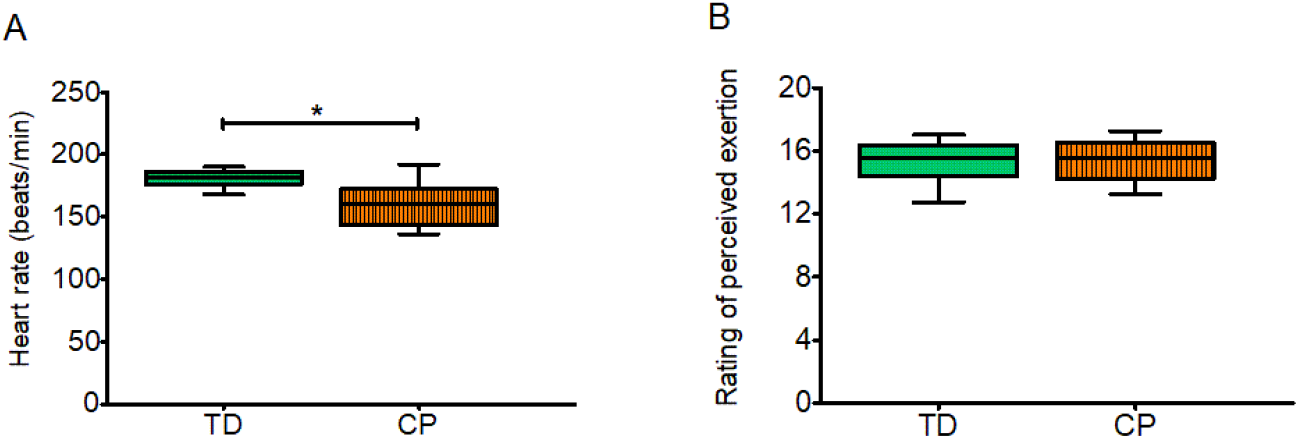
Mean heart rates (A) and ratings of perceived exertion (B) from the 45 min run in the participants with cerebral palsy (CP) and the typically developed participants (TD). The line in the middle of the boxes is plotted at the median, while the boxes extend from the 25th to 75th percentiles. Whiskers range from the smallest to the largest value. Significant difference: * p ≤ 0.001.

Regarding the baseline composition of the immune cell populations (Figure 3α and Figure 4α), a difference between groups was found in the TCRγδ+ T-cells (Figure 4Cα).

**Figure 3.**
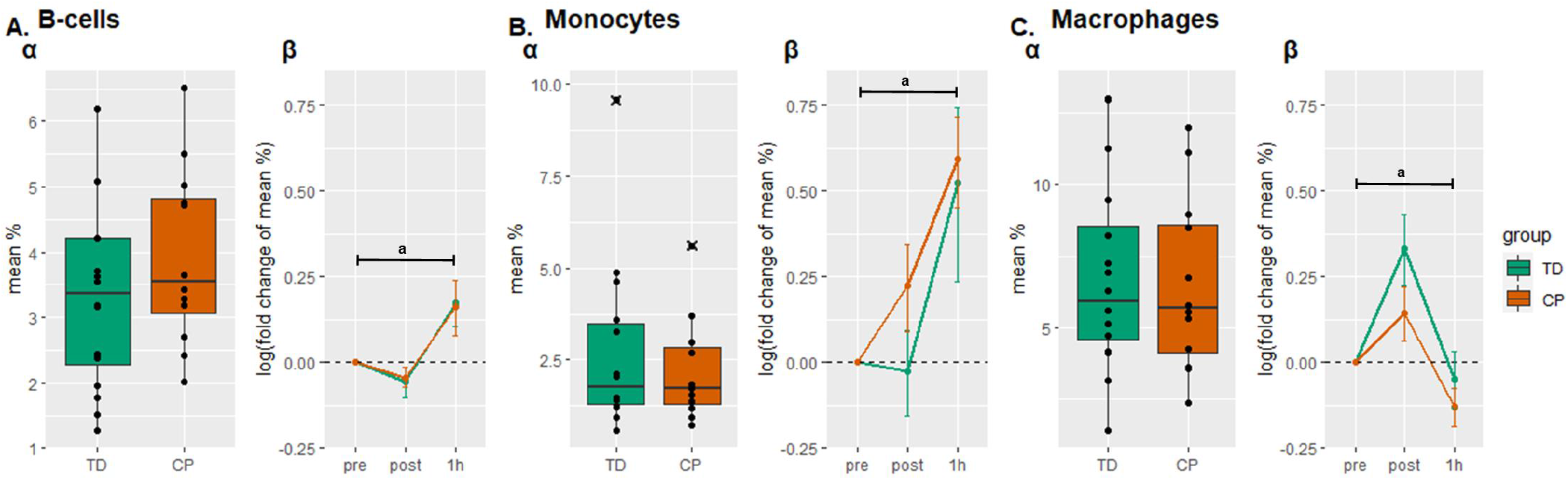
Myeloid panel. α) Baseline myeloid cell populations (% of total cells) of adults with cerebral palsy (CP) and typically developing (TD) adults. β) Changes of the cell populations immediately after (post) and one hour (1h) following a 45 minutes endurance exercise bout. Data is shown as fold change. A, B-cells; B, Monocytes; C, Macrophages. x, outliers. Values are means ± standard errors; a, main effect of time (p < 0.05).

**Figure 4.**
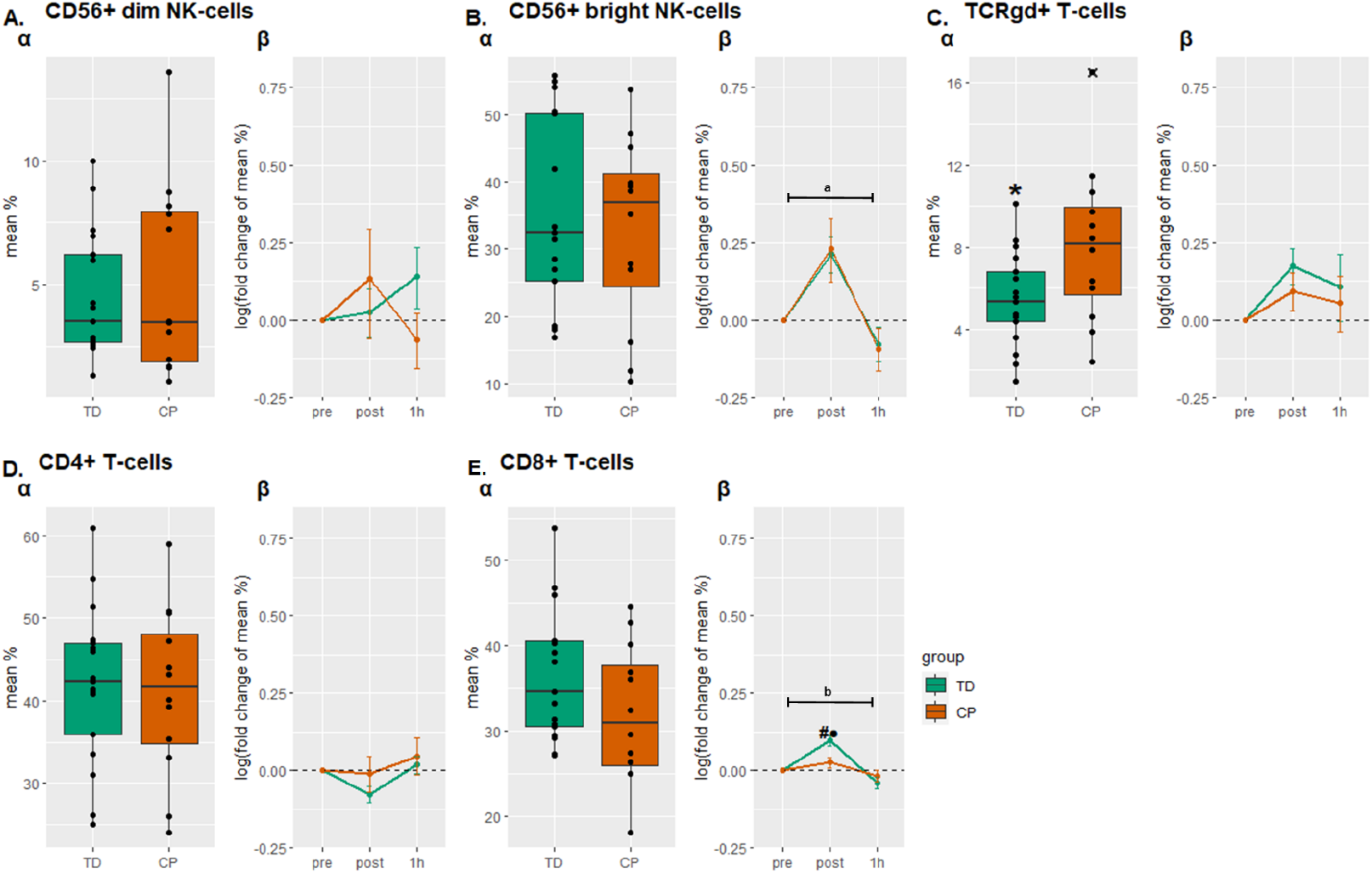
Lymphoid panel. α) Baseline lymphoid cell populations (% of total cells) for the adults with cerebral palsy (CP) and typically developing (TD) adults. β) Changes of the cell populations immediately after (post) and one hour (1h) following a 45-minute endurance exercise bout. Data is shown as fold change. A, CD56+ dim natural killer-(NK-) cells; B, CD56+ bright NK-cells; C, (TCRγδ+) T-cells; D, CD4+ T-cells; E, CD8+ T-cells; x, outliers. Values are as means ± standard errors; a, main effect of time (p ≤ 0.05); b, time*group interaction (p ≤ 0.05), # significant group difference at timepoint; • significantly different to pre in the TD individuals.

### Acute responses of circulating immune cell populations

In the myeloid panel, a significant main effect of time (p < 0.05) was found for the B-cells, monocytes, and macrophages (Figure 3A-C,β), indicating a general effect of exercise on these cell subpopulations in both individuals with CP and TD participants.

In the lymphoid panel, a significant effect of time (p < 0.05) was found in CD56+ bright NK-cells mainly due to an overall increase immediately following the acute exercise bout (Figure 4Bβ). Moreover, a significant time*group interaction (p < 0.05) was observed in the CD8+ T-cell population. The post-hoc comparison revealed that CD8+ T-cells increased after exercise in the TD participants but not in the individuals with CP (Figure 4Eβ). To evaluate the effect of exercise intensity on the response of the CD8+ T-cells, the response of these cells was additionally correlated to heart rate using a Pearson correlation. The analysis revealed a positive correlation between the two parameters (Figure 5; r = .537, p = .003).

**Figure 5.**
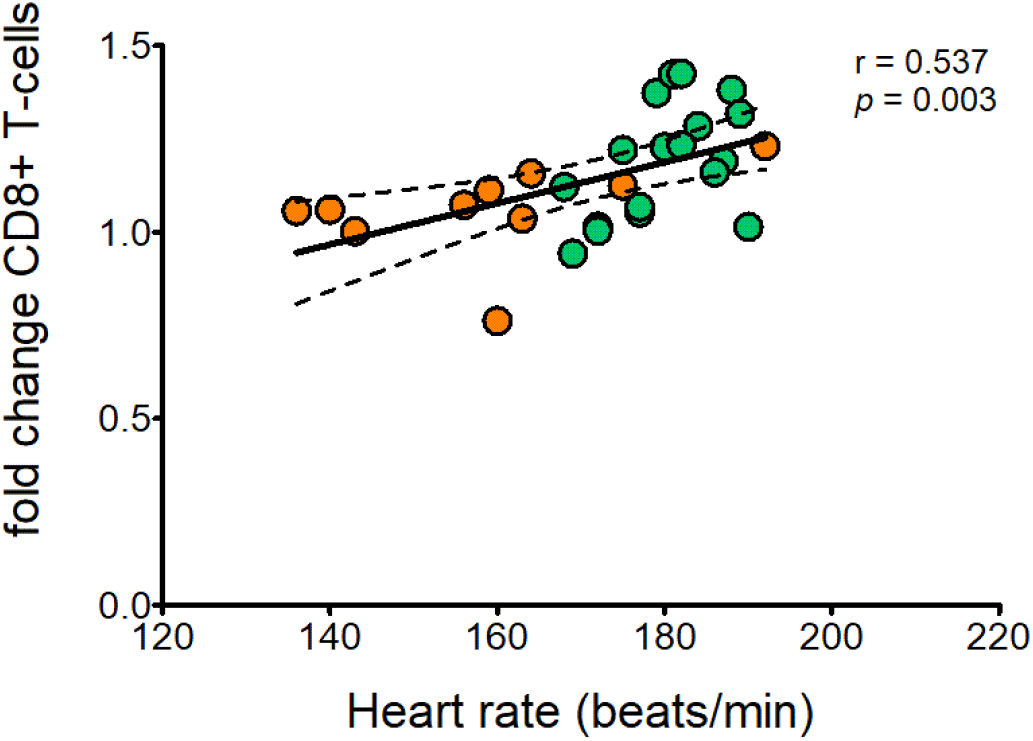
Positive correlation between mean heart rate (beats/min) and changes of the CD8+ T-cells population measured immediately following a 45-minute endurance exercise bout in individuals with cerebral palsy (orange circles) and their typically developing peers (green circles).

## Discussion

In this study, we assessed circulating immune cell populations in young adults with CP compared to their TD peers and examined the response of the cells to acute endurance exercise. The most important findings of the present study were that 1) at rest, the TCRγδ+ T-cell population was significantly different between the individuals with CP and TD, and 2) the CD8+ T-cells of the two groups responded differently to the acute endurance exercise bout, with significantly higher levels observed immediately after the exercise bout in the TD participants.

To date, little is known about the extent to which the immune system is involved in individuals with CP. For diseases of the central nervous system (CNS), studies emphasize the occurrence of chronic low-grade inflammation and immunodeficiency in patients with, for instance, spinal cord injury.^20^ In agreement, recent studies found markers of chronic low-grade inflammation also in young patients with CP. For instance, increased levels of TNF-α ^12,13^ and IL-6 levels ^13,21^ were observed in children with CP ≤ 5 years of age. Moreover, these studies indicate that the highest levels of pro-inflammatory cytokines are present in the youngest children (1-3 years).^12,21^ This finding is in line with the results presented by Pingel et al.^10^ and Zareen et al.^9^, who found no alterations in older children as well as adults with CP, when compared to healthy controls. Taken together, the mentioned studies support the relevance of age regarding the existence of pro-inflammatory cytokines/chronic inflammation in individuals with CP. Sharova and co-workers^13^ observed increased levels of CD3+ T-cells (+22%) and both CD4+ (+20%) and CD8+ T-cell (+21%) subsets along with increased levels of CD22+ B-cells (+28%) in young children with CP compared to TD children. In contrast, significant differences in most circulating immune cell populations could not be found between young adults with CP and TD in the current study. This result indicates a similar immune system state in individuals with CP and TD in adulthood. However, we found an increased baseline of TCRγδ T-cells in CP compared to TD. These rare T-cells connect innate and adaptive immune system, can exert strong anti-tumor cytotoxicity^22^, and have an extensive cytokine production capacity. Interestingly, they have previously been implicated in the immune systems response to periventricular leukomalacia in infants, a major cause of CP.^23^ In this context, TCRγδ T-cells appear to contribute to the injury of the brain by inducing amongst others the production of IL-17, subsequently leading to CNS inflammation and recruitment of other immune cell types that result in demyelinating lesions.^24^ While retained high levels of TCRγδ T-cells in adults have not been reported before, this may emphasize the continued relevance of this cell subtype into adulthood for its inflammatory and immune role in CNS disease. However, assessments of the cytokine profiles of young adults with CP are needed to clarify and to support this finding.

In addition to the quantification of the circulating immune cell populations at rest, we investigated the response of the circulating immune cell populations to an acute endurance exercise bout in both groups. While the cell responses observed in individuals with CP and TD are mostly in line with the behavior of the immune cells reported in previous studies,^7,16^ a different response was observed for the CD8+ T-cells. It is well established that endurance exercise leads to an increase in T-cells and their subsets, such as the cytotoxic CD8+ T-cells.^25,26^ In the present study, the participants with CP demonstrated a significantly lower number of circulating CD8+ T-cells directly after the acute bout of exercise compared with the TD participants. We assume that exercise intensity might have been the main influencing factor. The influence of exercise intensity on lymphocytosis has been previously described.^27^ Moreover, studies emphasize the intensity related response of the CD8+ T-cells to (endurance and resistance) exercise in typically developed people,^28,29^ with higher intensities causing elevated responses. In the present study, perceived exertion was rated as 15 in both groups, characterizing the endurance exercise bout as mostly “hard (heavy)”. Although no differences in perceived exertion were detected, the individuals with CP reached lower heart rates (−11.1%) in comparison to their TD peers. Since ratings of perceived exertion are subjective,^30^ heart rate data may be seen as more objective criteria to determine exercise intensity. Therefore, considering heart rate as an objective representation of the exercise-imposed strain on the cardiovascular system, this finding may suggest a lower strain evoked during the 45 minutes of Frame Running in the participants with CP. Since cardiovascular strain affects blood pressure, shear pressure in the blood vessels, and cardiac output, the lower intensity may have resulted in a lower level of immune cell mobilization and thus a lower number of CD8+ T-cells was measured from peripheral venous blood in this study. Supporting this assumption, we found a positive correlation between the response of the CD8+ T-cells to heart rate. The discrepancy between groups might have been cause by the performance of the endurance bout, i.e., Frame Running (cycling) vs. running, or the lower physical fitness level of the individuals with CP. Both aspects have to be further elucidated in future studies.

Notably, other factors such as poor motor control and motivation might also have had an influence on exercise performance and intensity, therefore also influencing the results of this study. Furthermore, the idea that the responsiveness of the CD8+ T-cells is reduced in general in individuals with CP cannot be verified. However, the responses of the other measured immune cells did not show significant alterations in comparison to those of the TD participants, which may indicate an altered function of CD8+ T-cells.

There are certain aspects that need to be considered when interpreting the results of the current study. Firstly, we note that the participating individuals with CP were already familiar with Frame Running and were mainly moderately physically active. Therefore, the findings might not be transferable to a less active population. Furthermore, the employed flow cytometric method of immune system analysis can only be a peripheral snapshot of immune system activity and should be interpreted as such. As we performed broad panel analysis rather than investigating individual sub-populations at a high resolution, there might also be additional shifts at the sub-population level that are not detectable with our analysis. Finally, investigation of cytokine profiles may be needed to verify our findings.

To the best of our knowledge, this is the first study investigating the immune status and immune cell responses in young adults with CP. In summary, no evidence for an impaired immune function could be found in this group when compared to their TD peers. The findings largely suggest a “recovery” of immune disturbances with age/maturation likely present throughout childhood. However, this should be further elucidated in future longitudinal studies. We further emphasize the importance of exercise intensity to evoke immune cell responses in individuals with CP. Thus, our data underline the importance of treatment and training strategies of adequate exercise intensities that cause positive responses of the immune system.

## Acknowledgments

The study was supported by grants of the Norrbacka-Eugenia Stiftelsen, Stiftelsen Sunnerdahls Handikappfond, and Stiftelsen Promobilia to FvW.

## Data availability statement

The data that support the findings of this study are available from the corresponding author upon reasonable request.

